# Respiration-Triggered Olfactory Stimulation Reduces Obstructive Sleep Apnea Severity – A Prospective Pilot Study

**DOI:** 10.1101/2023.02.28.530439

**Authors:** Ofer Perl, Lilach Kemer, Amit Green, Nissim Arish, Yael Corcos, Anat Arzi, Yaron Dagan

## Abstract

Obstructive sleep apnea (OSA) is a prevalent sleep-disordered breathing condition characterized by repetitive reduction in breathing during sleep. Current care standard for OSA is continuous positive air pressure devices, often suffering from low tolerance due to limited adherence. Capitalizing on the unique neurocircuitry of olfactory perception and its retained function during sleep, we conducted a pilot study to test transient, respiration-based olfactory stimulation as a treatment for OSA markers. Thirty-two OSA patients (Apnea-Hypopnea Index (AHI)≥15 events/hour) underwent two polysomnography sessions, ‘Odor’ and ‘Control’, in random order. In ‘Odor’ nights, patients were presented with transient respiratory-based olfactory stimulation delivered via a computer-controlled commercial olfactometer (Scentific). The olfactometer, equipped with a wireless monitoring, analyzed respiratory patterns and presented odor upon detection of respiratory events. No odors were presented in ‘Control’ nights. Following exclusions, 17 patients entered analysis (4 women, 47.4 (10.5) years, BMI: 33.8 (7.8)). We observed that olfactory stimulation during sleep reduced AHI (‘Odor’:17.2 (20.9), ‘Control’: 28.2 (18.6), z=- 3.337, p=0.000846, BF10=57.9), reflecting an average decrease of 31.3% in event number. Relatedly, stimulation reduced the oxygen desaturation index (ODI) by 26.9% (‘Odor’: 12.5 (15.8), ‘Control’: 25.7 (25.9), z=-3.337, p=0.000846, BF10=9.522. This effect was not linked to baseline OSA markers severity (ρ=-0.042, p=0.87). Olfactory stimulation did not arouse from sleep or affect sleep structure, measured as time per sleep stage (F(1,16)=0.088, p=0.77). In conclusion, olfactory stimulation during sleep was effective in reducing OSA markers severity without inducing arousals and may provide a novel treatment for OSA, prompting continued research.

## INTRODUCTION

Obstructive sleep apnea, a sleep-disordered breathing condition, is characterized by repetitive reductions in breathing while asleep due to collapse of the soft tissue of the upper airways^1^. OSA is an alarmingly prevalent disorder, with conservative estimates suggesting at least 7-9% of the general adult population are afflicted with OSA to some degree^2,3^.The impact of OSA on health and quality of life is widespread. OSA is linked with deleterious effects on daytime cognitive functions (e.g. attention^4^, memory^5^ and depression^6^), as well as increased morbidity and mortality as a result of accidents^7^ and long-term cardiovascular comorbidities^8,9^.

Continuous positive airway pressure (CPAP) treatment has long served as the primary treatment modality for OSA. Although effective, adherence to CPAP treatment regime is often suboptimal. To reach clinical efficiency, minimally acceptable levels of adherence to CPAP are at least four hours of treatment per day^10^. Estimates suggest only 30%–50% of patients reach this criterion and are often compelled to seek other treatment or go untreated^11^. Furthermore, chronic CPAP use is associated with side effects, such as nocturnal awakenings, oral dryness and naso-oral soreness^12,13^. While some alternatives to CPAP treatment exist^14^, such as oral appliances (e.g., mandibular advancement devices^15,16^), or more invasive procedures^17^, most are either less effective, come with various risks, or impose financial burden. As a result, CPAP remains the standard of care for efficient reduction of OSA markers.

In the search for an effective minimally-encumbering, noninvasive treatment for OSA, a central sensory pathway has been largely overlooked – olfactory stimulation during sleep. Olfactory stimulation possesses a unique combination of traits that are particularly attractive for OSA treatment: First, purely olfactory and mildly trigeminal odorants (trigeminality is defined as the ability to stimulate the branch of the trigeminal nerve nested within the olfactory epithelium^18^) do not wake from sleep^19^–^23^. Furthermore, evidence suggest sleep-benefiting effects for olfactory stimulation^24,25^ during sleep, in health^26^–^28^ and in clinical settings^29,30^. Second, and critically, despite not inducing arousal, incoming olfactory sensory information is picked up by the sleeping brain to influence physiology^19^, cognition^23,31^–^33^, and most importantly in the case of OSA – respiration^22,34^.

One core feature of olfactory processing is the ‘sniff response’, a sensorimotor loop by which presentation of odorants affects nasal airflow. This link between sensory input and respiration mode, established in wake humans^35,36^, is maintained during sleep^22,34,37^. Specifically, olfactory stimulation modulates sniff volume and duration in subsequent respiratory cycles during sleep. In previous experimentation, we were able to modulate respiratory patterns in healthy adult participants using odors during sleep^22^. Our findings suggested that transient olfactory stimuli modulated the inhalation/exhalation ratio during sleep. Thus, given that odors covertly manipulate respiration during sleep, it is conceivable to use olfactory stimulation to mitigate breathing disorders. A few studies have explored the link between olfaction and respiration in clinical settings, such as on apnea of prematurity in preterm neonates^38^–^40^, but not as a treatment for adult OSA. In addition, in most studies, odor delivery relied on transient in-room diffusers or a single acute exposure before sleep which are likely to decay due to sensory habituation^41^.

With this in mind we designed a pilot study to test the efficacy of transient, respiration-based olfactory stimulation as a treatment for OSA markers reduction. Owing to the fact that multiple experimental factors were tested in this design for the first time on OSA patients (e.g., olfactometer setup – odors and duration, odor delivery decision algorithm), this study was predefined to accommodate a limited number of participants and was therefore designated a pilot study. Specifically, we presented olfactory stimulation to OSA patients using a computer-control olfactometry device customized to fit individual needs. The olfactometer monitored patients’ respiratory patterns and presented the odors in a targeted manner upon detection of respiratory events.

## METHODS

### Patients

Thirty-two (7 women, 25 men, age 49.45 (11.18) years (median (IQR)) patients with a preexisting diagnosis of OSA with a severity threshold defined by an Apnea-Hypopnea Index (AHI) ≥ 15 events/hour polysomnography (PSG) were recruited and provided informed consent to participate in the study. Written informed consent to the procedures was approved by the ethics committees of Assuta Medical Center and Sha’are Zedek Medical Center.

A power analysis, computed on a preliminary pilot cohort (N = 5) suggested a decrease of 40%+/- 27.8 in AHI (Mean +/- SD). In light of these results we conducted a power analysis aimed at 5% alpha and 80% statistical power which revealed that a sample size of N = 20 could would a suitable cohort to demonstrate such effects. We predicted that ∼40% of participants could potentially be discarded, due to failure to complete the two-night course of the paradigm, difficulties falling asleep in the sleep clinic or AHI < 15 in the ‘control’ night due to variability in OSA severity over time. Thus, we opted to recruit a substantially larger cohort.

Inclusion and exclusion criteria are provided in Table 1. Following exclusions, the study cohort consisted of 17 patients (4 women, 13 men, age 47.4 (10.5) years, range: 30 – 66) patients. A CONSORT diagram detailing the dynamics of patient enrollment and inclusion is available in Supplementary Information. The demographics and clinical data of the 17 patients included in the study are detailed in Table 2.

**Table 1:** Inclusion and exclusion criteria for participation in the study.

| <u>Inclusion criteria</u> | <u>Exclusion criteria</u> |
| --- | --- |
| 1. OSA diagnosed; AHI≥15<br>2. Male and Female Aged 30 to 70 years old.<br>3. Normal Olfaction; Self-reported normal sense of smell.<br>4. Patients of childbearing potential must agree to use methods of contraception and have negative pregnancy test at screening. Effective methods of contraception must be used throughout the study.<br>5. BMI$<$ 35<br>6. Patient is willing and able to give his/her written informed consent. | 1. Chronic lung disease (including COPD, asthma).<br>2. Congestive Heart Failure.<br>3. Exhibiting any flu-like or upper respiratory illness symptoms at time of assessment.<br>4. History of severe nasal allergies or sinusitis or difficulty breathing through the nose.<br>5. Persistent blockage of one or both nostrils.<br>6. Any previous operation or trauma to the nose from the last 5 years.<br>7. Previous diagnosis of insomnia, narcolepsy, periodic limb movement disorder, respiratory failure.<br>8. Any use of antipsychotic, Hypnotic drugs.<br>9. Major neurological diagnosis.<br>10. Active malignant disease including chemotherapy or radiotherapy treatment.<br>11. Pregnant or lactating women.<br>12. Drug abuse.<br>13. Medical history of epilepsy.<br>14. C-PAP dependency; according to investigator discretion<br>15. COVID-19 diagnosed from the last 3 months.<br>16. Any known allergy to odorant capsules solution used during the study. |

**Table 2:**
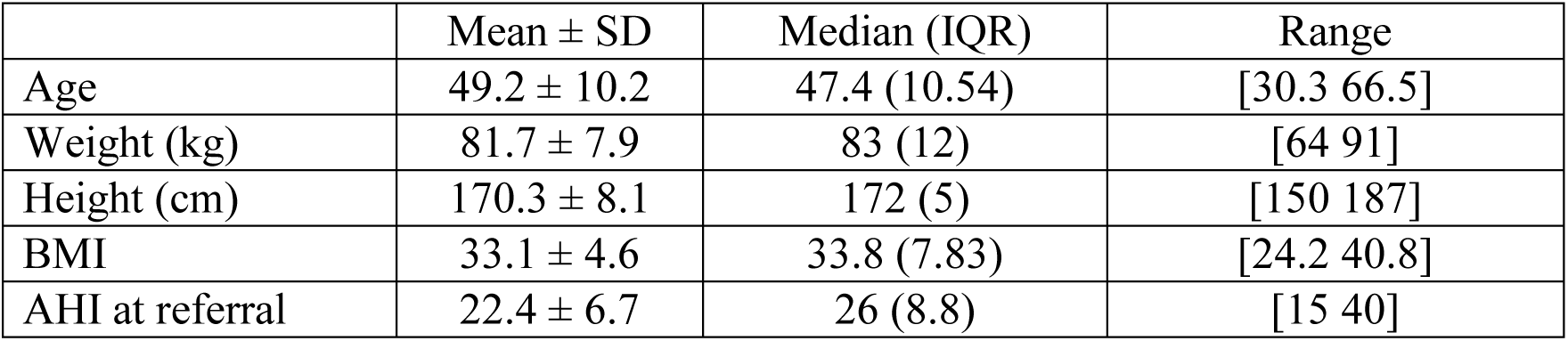
Baseline characteristics of the patient cohort. SD – standard deviation. IQR – interquartile range.

| | Mean ± SD | Median (IQR) | Range |
| --- | --- | --- | --- |
| Age | 49.2 ± 10.2 | 47.4 (10.54) | [30.3 66.5] |
| Weight (kg) | 81.7 ± 7.9 | 83 (12) | [64 91] |
| Height (cm) | 170.3 ± 8.1 | 172 (5) | [150 187] |
| BMI | 33.1 ± 4.6 | 33.8 (7.83) | [24.2 40.8] |
| AHI at referral | 22.4 ± 6.7 | 26 (8.8) | [15 40] |

### Study Design

This prospective, within-subject study was conducted at Assuta Medical Center, Tel-Aviv, and Sha’are Zedek Medical Center, Jerusalem, Israel, between October 2020 and September 2021. The study was registered in the online database clinicaltrials.gov (NCT04609618) and was approved by both medical centers’ ethics committees (E.C. #0045-20, E.C. #0560-20).

The study design comprised of two nights randomly ordered: treatment (‘Odor’) in which olfactory stimulation was delivered, and control (‘Control’), in which participants slept in the same setting but with no odor delivery, or any other OSA-related aid. The two conditions were presented in random order.

### OSA Severity Classification

The number of apneas and hypopneas per hour of sleep is collectively termed AHI. An apnea was defined as a reduction of > 90% in airflow and a hypopnea was defined as a reduction of 30-90**%** in airflow and/or effort, lasting > 10 seconds and accompanied by an arousal or a fall in SaO2 of >= 3%.

An AHI of 5 or more events per hour is a cutoff for OSA diagnosis. The AHI is further used to categorize OSA severity. Patients with AHI of 5 to 15, 16 to 30, or more than 30 events per hour, are considered to have mild, moderate, or severe OSA, respectively^43^. An additional marker used in OSA evaluation is the Oxygen Desaturation Index (ODI) defined as a 3% or more decrease in blood oxygen saturation (SpO2) below baseline. Notably, apnea severity and its clinical classification may change over time and in different sleeping conditions^42^–^44^. The time gap between OSA diagnosis and participation in the experiment was 337 (396) days in this cohort. Thus, in order to reevaluate participants’ current OSA severity, in a time window relevant for the experimental setting, we used the PSG measures recorded during the night without olfactory stimulation (‘Control’) as a baseline for the ‘Odor’ condition.

### Polysomnography

Polysomnography (PSG) was conducted and scored using the SOMNOscreen PSG platform (SOMNOmedics, Germany). The same recurring staff of sleep lab technicians in each center performed sleep data processing in accordance with the guidelines of American Academy of Sleep Medicine (AASM)^1^. PSG staff members were blind to the experimental conditions which were randomly assigned for each night. PSG montage consisted of a 6-lead electroencephalogram (EEG), two bilateral electrooculograms (EOG) leads, one submental lead, two tibial electromyograms (EMG), and one electrocardiogram (ECG) lead. Respiratory measurements in the PSG setup were evaluated from chest wall and abdominal movement using plethysmography as well as nasal pressure sensors and thermal airflow signals. Oxygen saturation and heart rate were monitored using a pulse oximeter. In addition, end-tidal carbon dioxide signals were obtained (EtCO2) (Respironics, USA) but not used during analysis due to poor signal quality. Lastly, sleep position, and audiovisual recordings were collected.

Processed data exported from the PSG platform included sleep continuity parameters, latency to stage 1 and stage, percent of time in wake after sleep onset (%WASO), index of awakenings, total sleep time (TST), and percent of time in bed (%TIB). Sleep architecture parameters consisted of percent of time spent in stage N1, stage N2, rapid eye movements (REM), and slow-wave sleep (SWS), index of sleep stage changes, and REM onset latency. Arousals were first labeled automatically by the PSG sleep scoring algorithm. Next, these labels were then verified and corrected if needed, by the sleep lab personnel (a technician and a physician).

### Olfactory stimulation

#### Odorants

All patients were presented with a fixed repertoire of five odor mixtures based on essential oils: orange, eucalyptus, lavender, frankincense-cinnamon and geranium (Phyto-Active, Israel). All odor materials held certificates of conformity with International Fragrance Association (IFRA) standards. Across the cohort, there was only a single case of a mild adverse event, feeling of nausea, that has been adjudicated as possibly related to the olfactory stimulation. No serious adverse events have been reported.

#### Olfactometer

Olfactory stimulation was carried out using a computerized olfactometer which is a commercial product designed to present odors at predefined times, for example in this study when an apnea event is detected (Scentific™, Appscent Medical). The olfactometer consisted of the following main components: a respiration detector, a controller, an odor disperser, and an odor cartridge (Fig. 1a). The respiration detector was equipped with a diagnostic system capable of online monitoring of patients’ respiratory patterns based on Impulse Radio Ultra-Wideband (IRUWB) radar^45–47^. The controller unit detected movement patterns associated with the onset of apnea-hypopnea epochs which then triggered odor delivery. Finally, the odor disperser consisted of a computer-controlled, and a low-pressure compressed air compartment in which disposable odorant cartridges were housed.

**Figure 1:**
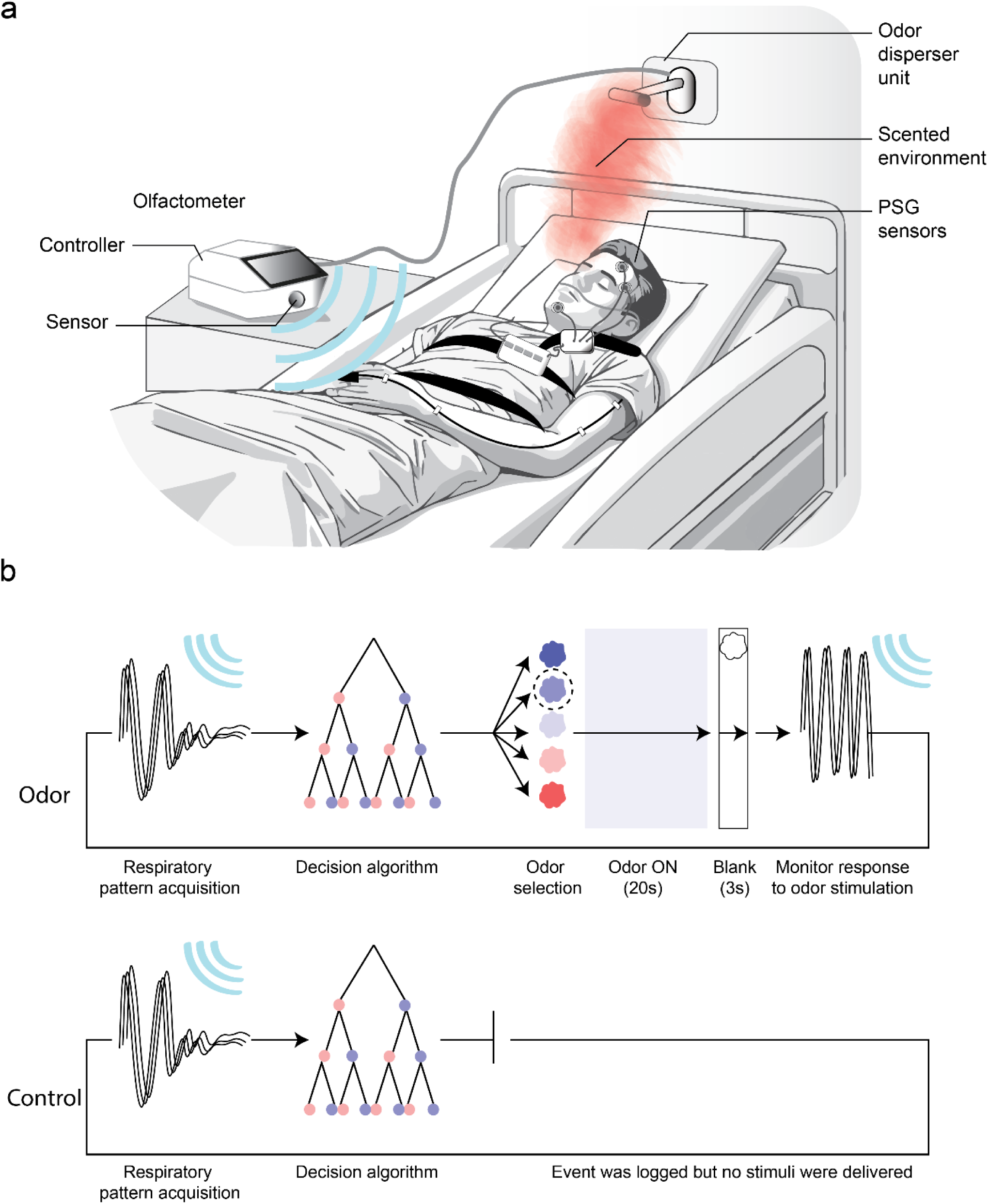
**(a)** Illustration of the experimental room. Patients spent two nights undergoing standard PSG. The olfactometer’s controller unit monitored and analyzed respiratory patterns using a contactless radio wave emitter. Olfactory stimulation was delivered via a wall-mounted disperser unit above the patient’s head. **(b)** Olfactory stimulation procedure. Top: in the ‘Odor’ condition respiratory patterns were continuously monitored and served as input to the decision algorithm. If characteristics of apnea- hypopnea patterns were met, the olfactometer dispersed one of five odorants selected at random for 20 seconds. Next, a 3-second burst of clean air (Blank) was presented. Finally, the olfactometer again monitored respiratory patterns for the next bout of this decision cycle. Bottom: in the ‘Control’ condition respiratory patterns were continuously monitored and served as input to the decision algorithm in order to log respiratory events, but no odors were dispersed. Note that contrary to the illustration, wireless monitoring did not stop during odor/blank delivery but persisted during odor presentation.

We positioned the odor disperser at about 50 cm from the patient’s head to target odor delivery at the vicinity of the nose. This setup allowed an odorous environment at the nose, devoid of other visual, tactile, humidity, or thermal stimuli.

#### Odor delivery regimen

The olfactometer constantly monitored respiratory patterns. When an apnea was automatically detected by the system algorithm, an odor was presented approximately one second later for a period of 20 seconds, concluding with an inter-stimulus-interval (ISI) of 40 seconds. Detected apnea events did not trigger odor presentation if they occurred within the ISI. The olfactometer rotated between five odors, presenting them in a random order to prevent olfactory habituation. Following each odor burst, non-odorous room air was emitted from the nozzles for three seconds to reduce odor contamination to the tubing (Fig. 1b). Apnea detection was based on a decision-tree algorithm, initially trained on 392 IRUWB signal features which were subsequently optimized to 20-feature repertoire. Critically, the olfactometer did not counteract apnea-hypopnea events only once they met the minimal duration of 10 seconds as clinically defined, but attempted to constantly predict their occurrence ahead of time. We first assessed the olfactometer’s apnea-hypopnea detection rate by comparing apnea-hypopnea events onsets between the olfactometer and the PSG, which underwent manual inspection by experienced sleep technicians. Since the olfactometer logged apnea-hypopnea events both in the ‘Odor’ condition and ‘Control’ (without applying the odor stimulation), we inspected both datasets.

### Data analysis and statistical approach

#### Statistical design

As a first step, the data underwent tests for normality using the Shapiro-Wilk test. We observed that the majority of parameters of interest, and specifically the OSA-related measures did not meet normality criterion (AHI: SW = 0.872, p = 0.0016; ODI: SW = 0.824, p = 0.0005). Therefore, we used non-parametric tests. Interactions were tested through a similar non-parametric approach, with tests aimed at contrasting the effects of two comparable measures in a 2-by-2 design (e.g. the interaction between effects of odor stimulation on apnea and hypopnea indices). All analyses described in this study were conducted using two-tailed tests. Analyses were carried out using custom code in MATLAB R2018a (MathWorks, Natick, MA) and JASP 2019 version 0.16.4.

Additionally, we included Bayesian non-parametric paired tests where appropriate, in order to provide additional information regarding the strength of evidence in favor of the null hypothesis^48^. Specifically, we compared the probability that odor presentation had no effect on a parameter (H0), to the alternate hypothesis that this parameter will change during the ‘Odor’ condition (H1) with a Cauchy prior of 0.707^49^. In line with the frequentist hypothesis testing described above, Bayesian tests of effects of olfactory stimulation on OSA also relied on two-sided assumptions. All Bayesian statistical analyses were conducted in JASP 2019 version 0.16.4.

## RESULTS

### Olfactory stimulation during sleep reduced apnea-hypopnea events in OSA patients

First, we set out to test the hypothesis that olfactory stimulation during sleep decreased OSA markers. We observed that in nights in which olfactory stimulation was delivered, AHI indices were significantly reduced (AHI ‘Odor’: 17.2 (20.9), AHI ‘Control’: 28.2 (18.6), Wilcoxon signed-rank test (14/17 patients): z = -3.337, p = 0.000846, BF10 = 57.9), reflecting an average decrease of 31.3% in the number of events (Fig. 2a). We then set out to ask whether olfactory stimulation had a sleep stage-specific beneficial effects on AHI. To this end we calculated sleep stage-specific AHI (the frequency of apnea-hypopnea events in each sleep stage) and contrasted these results between ‘Odor’ and ‘Control’ nights. We observed that when no olfactory stimulation was applied, apnea-hypopnea events occurred predominantly in N2 sleep (AHI = 33.5 (18.1)) and REM sleep (AHI =29.1 (22.9)) and less frequently in N3 sleep (AHI = 5.3 (30.2). When odor stimulation was applied, the stage-specific AHI was significantly reduced during N2 sleep (‘Odor’: 16.28 (32.4), ‘Control’:33.54 (18.1), Wilcoxon signed-rank: z = -2.34, p = 0.019, BF10 = 2.47), and N3 sleep (‘Odor’: 2.26 (4.7), ‘Control’: 5.36 (30.23), Wilcoxon signed-rank: z = -2.79, p = 0.032, BF10 = 2.08). The same trend was observed during REM sleep but it did not reach significance (‘Odor’: 21.73 (26.4), ‘Control: 29.06 (22.9), Wilcoxon signed-rank: z = -1.63, p = 0.102, BF10 = 0.80). Finally, we compared the effects across sleep stages by directly contrasting odor-mediated difference ([‘Odor’ – ‘Control’]) and observed that effects in NREM sleep were not significantly higher than REM sleep (‘N2’ vs. ‘REM’ Wilcoxon signed-rank: z = -0.83, p = 0.431, BF10 = 0.31; ‘N3’ vs. ‘REM’ Wilcoxon signed-rank: z = -0.64, p = 0.548, BF10 = 0.35). These results suggest that olfactory stimulation effectiveness in reducing AHI is sleep-stage independent.

**Figure 2:**
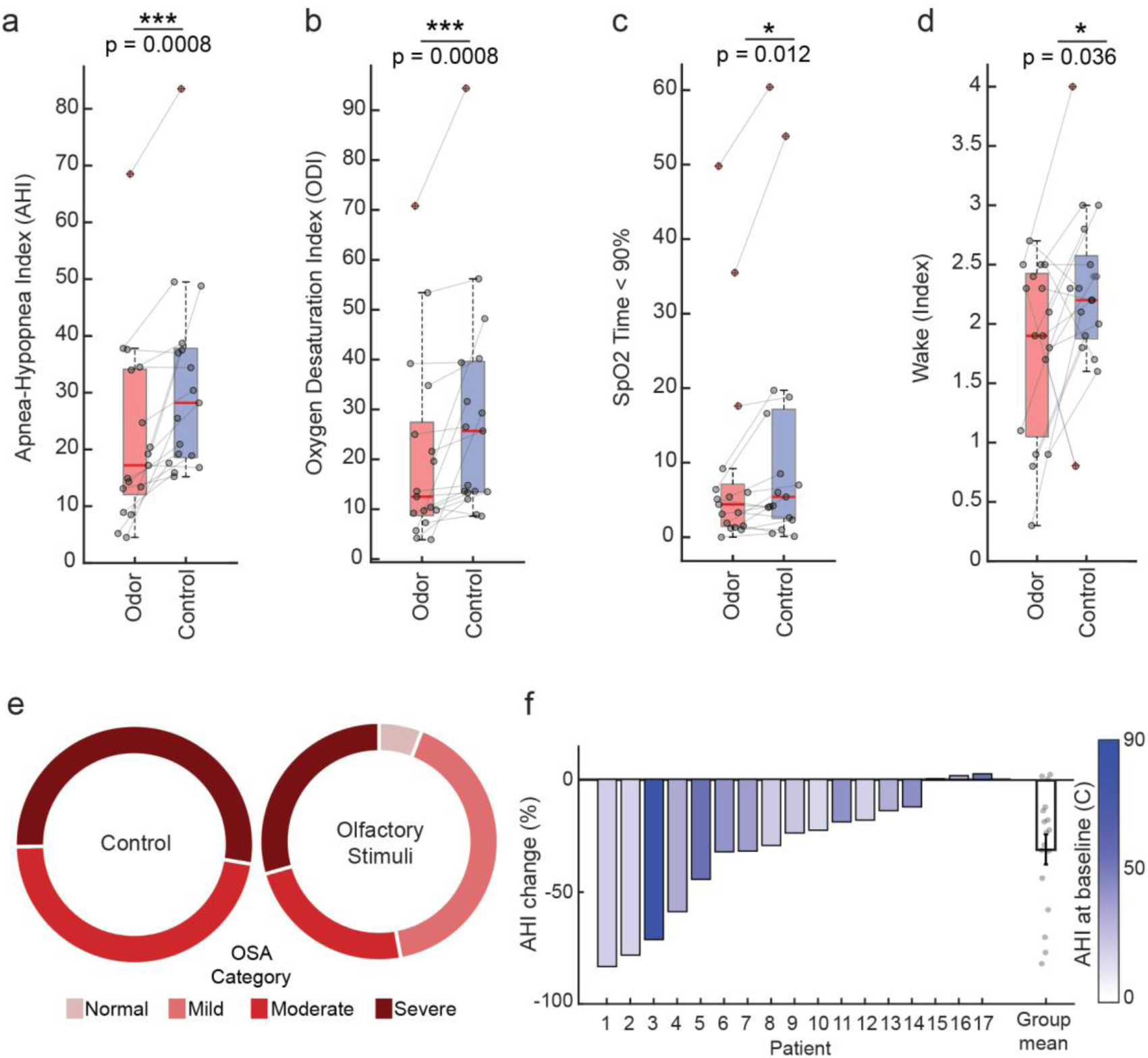
**(a)** Box plots depicting median-based group distribution of Apnea-Hypopnea Index (AHI) scores for ‘Odor’ and ‘Control’ nights (red and blue respectively). Points represent participants (N=17), with cross-condition lines overlaid to aid in visualization of within-subject effects. Thick red bars denote group median. Dashed whiskers extend to 25^th^ and 75^th^ percentile. Data points exceeding this range are marked with red crosses. * p < 0.05,*** p < 0.001, **(b)** Same as **(a)** but for the effect of olfactory stimulation on measures of Oxygen Desaturation Index (ODI) **(c)** Same as **(a)** but for the effect of olfactory stimulation on the percentage of time spent in oxygen saturation (SPO2) lower than 90%. **(d)** Same as **(a)** but for the effect of olfactory stimulation on Wake index. **(e)** Categorical change in apnea severity classification following olfactory stimulation during sleep (N=17). A breakdown of OSA classification in the ‘Control’ night (left) versus when ‘Olfactory stimuli’ were presented (right). Color intensity signifies OSA severity categories ranging from normal (light red) to severe (dark red). **(f)** Bars depicting individual (%) change of Apnea-Hypopnea Index in ‘Odor’ nights compared to ‘Control’ nights. Color map denotes OSA severity according baseline AHI measured in the ‘Control’ night. Rightmost bar depicts (%) change of the group mean. Points represent participants (N=17). Error bars are SEM.

Next, we confirmed that the chronological order of conditions had no effect on changes in OSA markers. We employed the same analytical approach, this time modeling session order (‘Night 1’ / ‘Night 2’) instead of conditions of olfactory stimulation. The two sessions took place 7 (3) days apart (Note that time between sessions was calculated with the exclusion of one patient who had to be summoned for a third session which took place 177 days following their previous visit, due to a technical issue with data recording in one of the nights). OSA-related respiratory measures were not affected by session order (AHI: Night 1: 24.7 (22.1) Night 2: 20.4 (16.8), Wilcoxon signed-rank test (8/17 patients): z = 0.40237, p = 0.68740, BF10 = 0.32), suggesting that the observed reduction in AHI following odor stimulation during sleep is not a result of conditions order.

Together, these findings demonstrate that transient odor stimulation during sleep decreases AHI in OSA patients.

Following odor presentation, patients’ OSA severity clinical categories was either improved or remained stable as follows: **Moderate OSA**: Out of the nine patients, seven improved: one (11.1%) to normal AHI (< 5 events/hour) and six (66.6%) to mild OSA classification while two (22.2%) patients remained in the moderate category. **Severe OSA**: Out of seven patients, three improved: one (12.5%) patient improved to mild OSA, and two (25%) improved to moderate OSA while five (62.5%) patients remained in the severe OSA category (Fig. 2e). Importantly, in none of the patients OSA severity according to clinical categories exacerbated. We observed between-patient variability in the magnitude of AHI reductions following odor stimulation (Fig. 2f) and therefore asked whether this variability is linked to baseline OSA markers severity (i.e., AHI in ‘Control’ = 28.2 (18.6), range: 15.2 – 83.5). Nevertheless, these two measures were not correlated (Spearman’s ρ = -0.042, p = 0.87).

In line with the improvement in AHI, we observed a reduction in oxygen desaturation index (ODI) and measurements related to oxygen saturation (SpO2). Olfactory stimulation reduced ODI by 26.9 % on average (ODI ‘Odor’:12.5 (15.8), ODI ‘Control’: 25.7 (25.9), Wilcoxon signed-rank test (16/17 patients): z = -3.3373, p = 0.000846, BF10 = 9.522, Fig. 2g) and reduced the proportion of time spent at SpO2 < 90% (‘Odor’: 4.4 (4.9), ‘Control’: 5.4 (14.0) %, Wilcoxon signed-rank test: z = -2.51, p = 0.0121, BF10 = 3.966, Fig 2c) In addition, marginal reduction in the average depth of SpO2 desaturation was observed (‘Odor’: 5.688 ± 1.369, ‘Control’: 6.424 ± 2.487, Wilcoxon signed-rank test: z = -1.94, p = 0.0517, BF10 = 1.586). Mean and minimal SpO2 were not significantly altered by olfactory stimulation (Mean SpO2: ‘Odor’: 93 (1), ‘Control’: 93 (2), Wilcoxon signed-rank test: z = -1.68, p = 0.125, BF10 = 0.832; Minimal SpO2: ‘Odor’: 84 (4), ‘Control’: 82 (5), Wilcoxon signed-rank test: z = 1.13, p = 0.271, BF10 = 0.451). Please refer to Table 3 for a summary of the results reported in this section. Together, these findings suggest that odor presentation during sleep reduces the severity of respiratory OSA-related markers.

**Table 3:** Effects of olfactory stimulation on OSA-related parameters: summary of the results.

| **Measurement** | **Odor** | **Control** | **z** | **p** |  | **BF<sub>10</sub>** |
| --- | --- | --- | --- | --- | --- | --- |
| Session duration (hr) | 6.78 (0.85) | 6.95 (1.23) | -1.396 | 0.174 | - | 0.763 |
| AHI (total) | 17.2 (20.9) | 28.2 (18.6) | -3.34 | 0.000846 | *** | 57.9 |
| ODI | 12.5 (15.8) | 25.7 (25.9) | -3.337 | 0.000846 | *** | 9.52 |
| Mean SpO<sub>2</sub>(%) | 93 (1) | 93 (2) | -1.668 | 0.125 | - | 0.832 |
| Minimal SpO<sub>2</sub> (%) | 84 (4) | 82 (5) | 1.13 | 0.271 | - | 0.451 |
| Time at SpO<sub>2</sub><90% (%) | 4.4 (4.9) | 5.4 (14.0) | -2.51 | 0.0121 | * | 3.966 |
| Event duration (s) | 20 (4) | 20 (4.5) | -1.16 | 0.263 | - | 0.26 |
| Wake (index) | 1.9 (1.37) | 2.2 (0.7) | -2.09 | 0.036 | * | 1.958 |
| Arousals (index) | 32.9 (26.12) | 26.9 (16.05) | -1.585 | 0.113 | - | 0.855 |
| **Measurement** | **Night 1** | **Night 2** | **z** | **p** |  | **BF<sub>10</sub>** |
| AHI (effect of order) | 24.7 (22.1) | 20.4 (16.8) | 0.402 | 0.687 | - | 0.32 |
| %TIB at wake (effect of order) | 13.2 (8) | 8.6 (5.6) | 2.627 | 0.0086 | ** | 6.456 |
| Sleep efficiency (effect of order) | 86.8 (8) | 91.4 (7.7) | 2.627 | 0.0086 | ** | 5.797 |

### Olfactory stimulation during sleep did not alter apnea-hypopnea events duration in OSA patients

Next, we tested whether olfactory stimulation during sleep reduced the duration of apnea-hypopnea events. First, we examined all five odors combined and found no difference in event duration between ‘Odor’ and ‘Control’ nights (‘Odor’: 20 (4) s, ‘Control’: 20 (4.5) s, Wilcoxon signed-rank test (12/17 patients): z = -1.160, p = 0.263, BF10 = 0.260). To examine the effect of respiration- based transient odor stimulation, we compared the duration of apnea-hypopnea events in the ‘Odor’ nights, that were conjugated with odor stimulation (i.e., the respiratory event which was the trigger for odor delivery) and apnea-hypopnea events that were not conjugated with odor stimulation. Overall the olfactometer presented 103 (66) odor pulses per treatment night. The olfactometer’s detection rate for events marked by the PSG was 50.35 (24.26) %, and did not differ between ‘Odor‘ and ‘Control’ nights (‘Odor’: 46.74 (20.04) %, ‘Control’: 50.38 (28.61) %, Wilcoxon signed-rank: z = -0.213, p = 0.854, BF10 = 0.250). In other words, approximately half of the apnea-hypopnea events were conjugated with odor stimulation.

We found no difference in apnea-hypopnea event duration between the two conditions (‘Conjunction: 19.5 (5.3) s, ‘Non-Conjunction’: 20.5 (5.18) s, Wilcoxon rank sum test (11/17 patients): z = -0.828, p = 0.431, BF10 = 0.343). Then, we examined the influence of each odor separately. Over the course of the night, olfactory stimulation was delivered using a fixed repertoire of five odor mixtures, primarily in order to reduce potential sensory habituation to recurrent exposures. To test whether some odors were consistently more effective than others in reducing OSA-related markers we sampled the duration of apnea-hypopnea events that were conjugated with each odor type. Normalized apnea-hypopnea events duration (within patient using z-score) showed no consistent effect of odorant type at the group level (rmANOVA F(4,64) = 0.5891, p = 0.672). Given known individual variability in odor perception ^52,53^, we repeated the analysis at the single-patient level. Again, we observed no differences in conjugated apnea-hypopnea events as a function of odor type. These findings imply that odor stimulation during sleep does not reduce the duration of apnea-hypopnea events.

### Olfactory stimulation did not promote arousal or alter sleep structure

Sleep quality questionnaires obtained during screening confirmed that the OSA patient cohort, on average, reported poor sleep quality. The median Pittsburgh Sleep Quality Index (PSQI) score was 4.5 (3) (N = 12), with a score of five typically serving as criterion for effectively distinguishing good and poor sleeper^50^. Similarly, median Epworth Sleepiness Scale (ESS) score was 10 (9) which is indicative of higher-normal daytime sleepiness^51^.

It is largely held that olfactory stimulation does not perturb sleep, and as discussed earlier, may promote it^24^–^30^. Nevertheless, in order to validate that in our experimental setting the olfactory stimulation did not arise from sleep, we compared the wake and arousal indices of ‘Odor’ and ‘Control’ nights. We found that ‘Odor’ nights had lower wake index than ‘Control’ nights (‘Odor’: 1.9 (1.37), ‘Control’: 2.2 (0.7), Wilcoxon signed-rank test (12/17 patients): z = -2.0956, p = 0.0361, BF10 = 1.958, Fig 2d), and arousal scores (both cortical and autonomic), were not significantly different between the two nights (Arousal total sleep: ‘Odor’: 32.9 (26.12), ‘Control’: 26.9 (16.05), Wilcoxon signed-rank test (12/17 patients): z = -1.585, p = 0.113, BF10 = 0.855). These findings indicate that odor stimulation did not induce arousals and, in line with the literature^25^ reduced the number of awakenings.

We then asked whether olfactory stimulation altered sleep structure, quantified as percent of time in bed (%TIB) spent in wakefulness and in each sleep stage (Table 4). A repeated measures ANOVA with conditions of ‘Stimulation (Odor/Control), and ‘Stage (Wake / REM / N1 / N2 / N3) did not uncover a main effect of ‘Stimulation’ (F(1,16) = 0.088, p = 0.77), implying no change in the time spent in each stage between the two nights. In addition, as expected we observed that on the first night in the sleep lab, regardless of whether odors were presented, the %TIB spent at wake was significantly higher (%TIB at wake: ‘Night 1’:13.2 (8), ‘Night 2’: 8.6 (5.6), Wilcoxon signed-rank test (11/17 patients): z = 2.6273, p = 0.00860, BF10 = 6.456) and sleep efficiency was lower (sleep efficiency: ‘Night 1’: 86.8 (8.0), ‘Night 2’: 91.4 (7.7), Wilcoxon signed-rank test (11/17 patients): z = 2.6273, p = 0.00860, BF10 = 5.797). Both these factors can be attributed to ‘first night’ effects due to sleeping in the unfamiliar lab setting^54^. In line with previous studies^19^–^23^, these findings demonstrate that odor presentation during sleep does not disturb sleep in OSA patients or alters sleep architecture significantly.

**Table 4:** Sleep structure distribution provided as % time spent in bed (%TIB), as well as arousals and wake index. Values are listed for ‘Odor’ and ‘Control’, per patient and sleep stage, separated by a vertical line. The bottom lines provide a summary of each field as Median (IQR) and the result of a statistical test contrasting between ‘Odor’ and ‘Control’ values.

| <u>Patient</u> | <u>Wake</u><br><u>(%) TIB</u> | <u>REM</u><br><u>(%) TIB</u> | <u>N1</u><br><u>(%) TIB</u> | <u>N2</u><br><u>(%) TIB</u> | <u>N3</u><br><u>(%) TIB</u> | <u>Arousals</u> | <u>Wake</u><br><u>(index)</u> |
| --- | --- | --- | --- | --- | --- | --- | --- |
|  | <u>Odor \|</u><br><u>Control</u> | <u>Odor \|</u><br><u>Control</u> | <u>Odor \|</u><br><u>Control</u> | <u>Odor \|</u><br><u>Control</u> | <u>Odor \|</u><br><u>Control</u> | <u>Odor \|</u><br><u>Control</u> | <u>Odor \|</u><br><u>Control</u> |
| 1 | 7.9 \| 7.8 | 17.8 \| 15.6 | 0.6 \| 1.1 | 42.5 \| 44.2 | 30.7 \| 31.3 | 14.2 \| 17.7 | 0.9 \| 1.9 |
| 2 | 16.4 \| 12 | 2.2 \| 15.8 | 2.5 \| 2.2 | 68.3 \| 61.3 | 10.6 \| 8.7 | 8.9 \| 22.2 | 2.5 \| 1.7 |
| 3 | 8.6 \| 10.6 | 8.8 \| 13 | 1.8 \| 0.5 | 75.6 \| 65.9 | 5.1 \| 10 | 22.5 \| 92.1 | 1.9 \| 2.2 |
| 4 | 17.8 \| 9 | 12.5 \| 1 | 0.4 \| 0.6 | 48 \| 82 | 21.3 \| 7.3 | 50.1 \| 44.2 | 1.8 \| 2.0 |
| 5 | 14.7 \| 18.1 | 12 \| 8.2 | 2.3 \| 0 | 58.5 \| 56.1 | 12.4 \| 17.6 | 33.9 \| 36.9 | 1.7 \| 2.4 |
| 6 | 5.9 \| 6.7 | 12.2 \| 17.5 | 0.5 \| 1.3 | 58.4 \| 58.4 | 23 \| 16.1 | 24.3 \| 3.9 | 2.5 \| 2.5 |
| 7 | 13 \| 4.9 | 21.2 \| 25 | 0.4 \| 0.5 | 46.8 \| 50.7 | 18.6 \| 19 | 38.8 \| 57.2 | 2.7 \| 2.3 |
| 8 | 16.8 \| 14.9 | 17.6 \| 20.3 | 0.5 \| 2 | 47 \| 52.4 | 18.1 \| 10.4 | 51.1 \| 32.4 | 0.3 \| 2.1 |
| 9 | 1.5 \| 7 | 17.1 \| 24.8 | 0.5 \| 0.3 | 50.3 \| 43.7 | 30.6 \| 24.2 | 52.9 \| 53.1 | 0.9 \| 1.6 |
| 10 | 7.6 \| 9.8 | 6.6 \| 9.7 | 1 \| 0.6 | 58.8 \| 49.4 | 26 \| 30.3 | 47.3 \| 35.5 | 2.4 \| 3.2 |
| 11 | 26.8 \| 43.2 | 10.9 \| 10.3 | 4.6 \| 5.3 | 39.1 \| 36.8 | 18.5 \| 3.5 | 23.7 \| 45.5 | 2.5 \| 4.0 |
| 12 | 6.4 \| 16.1 | 15.4 \| 10.3 | 3.1 \| 5.9 | 44 \| 51.1 | 23.1 \| 16.6 | 38.4 \| 47.3 | 2.1 \| 3.0 |
| 13 | 12.5 \| 13.2 | 18.2 \| 14.6 | 6.3 \| 4.7 | 44.8 \| 54.1 | 18.2 \| 13.4 | 13.6 \| 50.9 | 1.9 \| 2.8 |
| 14 | 22.4 \| 8.6 | 8.6 \| 5.6 | 11.5 \| 5.1 | 51 \| 58.9 | 6.6 \| 15.4 | 44 \| 43.6 | 2.3 \| 0.8 |
| 15 | 16.5 \| 6.9 | 8.3 \| 7.4 | 4.9 \| 7.5 | 59.2 \| 59.7 | 9.9 \| 18.6 | 32.9 \| 34.8 | 1.1 \| 2.3 |
| 16 | 20 \| 9.2 | 7.7 \| 8.2 | 7.8 \| 7.4 | 49.3 \| 60.2 | 15.1 \| 14.8 | 13.9 \| 31.4 | 2.3 \| 2.2 |
| 17 | 2.8 \| 2.3 | 10.2 \| 15.9 | 1.5 \| 1.2 | 68.6 \| 59.7 | 16.1 \| 20.9 | 20.2 \| 36.0 | 0.8 \| 1.8 |
| Median<br>(IQR) | 13 (9.2) \|<br>9.2 (6.2) | 12 (8.5) \|<br>13 (7.7) | 1.8 (4.1) \|<br>1.3 (4.5) | 50.3 (12) \|<br>56.1 (9) | 18.2 (10.6)<br>\| 16.1 (8.6) | 32.9 (23.8) \|<br>36.9 (14.9) | 1.9 (1.3) \|<br>2.2 (0.6) |
| p value | 0.844 | 0.977 | 0.836 | 0.956 | 0.402 | 0.113 | 0.036 |

## DISCUSSION

In this pilot study, we asked whether olfactory stimulation during sleep can alleviate sleep apnea markers. We leveraged on the unique ability of odors to exert physiological and neurological effects during sleep without inducing arousal^23,33^. Our main finding is that transient olfactory stimulation during sleep was associated with a pronounced reduction in patients’ AHI. Out of a cohort of 17 patients, 14 patients (82%) exhibited a reduction in AHI while the remaining three (18%) showed very subtle increases (all < 1 AHI). AHI reductions were not related to baseline OSA markers severity in this cohort (15 < AHI < 83 at baseline). To our knowledge, this is the first study to report beneficial effects of olfactory stimulation during sleep on OSA in adults.

Our study diverges from previous uses of olfactory stimulation in sleep in two central aspects. First, odors were delivered in transient pulses. Second, we used a rotating selection of odors rather than adhering to one. Both these aspects may have led to less sensory habituation compared to methods of constant passive exposure^55,56^. These improvements were achieved thanks to recent technological advancements in conditionally-triggered olfactometry^37,57^, miniaturization^58^, and on-the-fly processing with machine learning. Compared to the manually-validated PSG recording, the olfactometer detected about half of the apnea-hypopnea events detected by PSG. Notably, even though approximately half of events went undetected, we observed a reduction of over 30% in AHI. We speculate that higher detection rates may result in further reduction in AHI. That said, the dynamics of the respiratory response to odors are unclear and a cumulative effect may be exerted by sparse odor presentations as well.

OSA severity is linked to olfactory dysfunction^59,60^ yet the exact pathophysiological reasons are unclear^61^. During screening patients reported subjectively normal olfaction but did not undergo an objective evaluation (e.g., ‘Sniffin’ Sticks^62^). We speculate the patients with an intact sense of smell will be optimal candidates for this treatment method, given that severe olfactory dysfunction could hinder the perception of incoming sensory stimulation. Follow-up studies should dwell on this point in order to systematically delineate the precise relationship between olfactory abilities and the influence of olfactory stimulation during sleep on OSA.

Our findings complement evidence from research on neonatal apnea of prematurity^38,39^ (but see^63^ for opposite results) who reported ambient exposure to vanillin reduced apnea symptomatology. Here, odor identity did not affect event duration and no single odor type emerged as most effective. This observation aligns with previous studies showing that during sleep odor stimulation bears effects regardless of its identity or valence^23,33,64^. Future studies may systematically test different odorants to tackle this question directly. Olfactory stimulation did not alter sleep structure. However, we cannot disentangle the factors contributing to sleep continuity – the reduction in apneic events and the sleep-promoting effects of odors. We did not observe olfactory-induced improvements in parameters of sleep quality (e.g., %SWS, sleep efficiency).

The neurological mechanisms at play in OSA are unclear. Several characteristics of OSA suggest involvement of the central nervous system in the genesis or maintenance of the syndrome^65^. For example, obstruction in OSA is frequently preceded by short breathing pauses, apparently of central origin^8^. Which neurological mechanism may mediate the effects of olfactory stimulation on respiration? We propose that the mechanism by which breathing is reinstated is through the aspiration reflex. Nasopharyngeal stimulation was shown to restore spontaneous breathing in rodents and cats^66,67^, most likely through medullary nuclei of the pre-Bötzinger complex, a respiratory pace-maker^68^. Olfactory stimulation may act as a low-amplitude form of nasopharyngeal stimulation, thereby inducing the aspiration reflex without arousal. In line with this proposed mechanism, olfactory stimulation can potentially offer significant relief from central sleep apnea but this remains to be tested empirically.

Finally, we would like to highlight several limitations of our study. First, due to multiple exclusions, our sample size consisted of 17 patients, which limits the generalizability of the observed effects, and may have obscured additional effects due to low statistical power. Second, our results should be replicated in a more longitudinal design involving multiple nights per condition in order to test gradual cumulative effects and reduce naturally occurring between-night variability in OSA measures. Third, no reports of subjective outcome were collected and this should be rectified in future studies. Lastly, the low number of females in the cohort prevented us from testing any gender-related differences.

In conclusion, the high prevalence of OSA combined with current insufficient solutions warrant the search for alternative treatments a high priority. Our results suggest that contactless transient respiratory-based olfactory stimulation during sleep is a viable such alternative, prompting continued research on this front.

## Acknowledgments

We thank Dr. Arie Oksenberg for his insightful comments.

## Data Availability

The data underlying this article will be shared on reasonable request to the corresponding author.

---

**Supplementary Figure 1:**
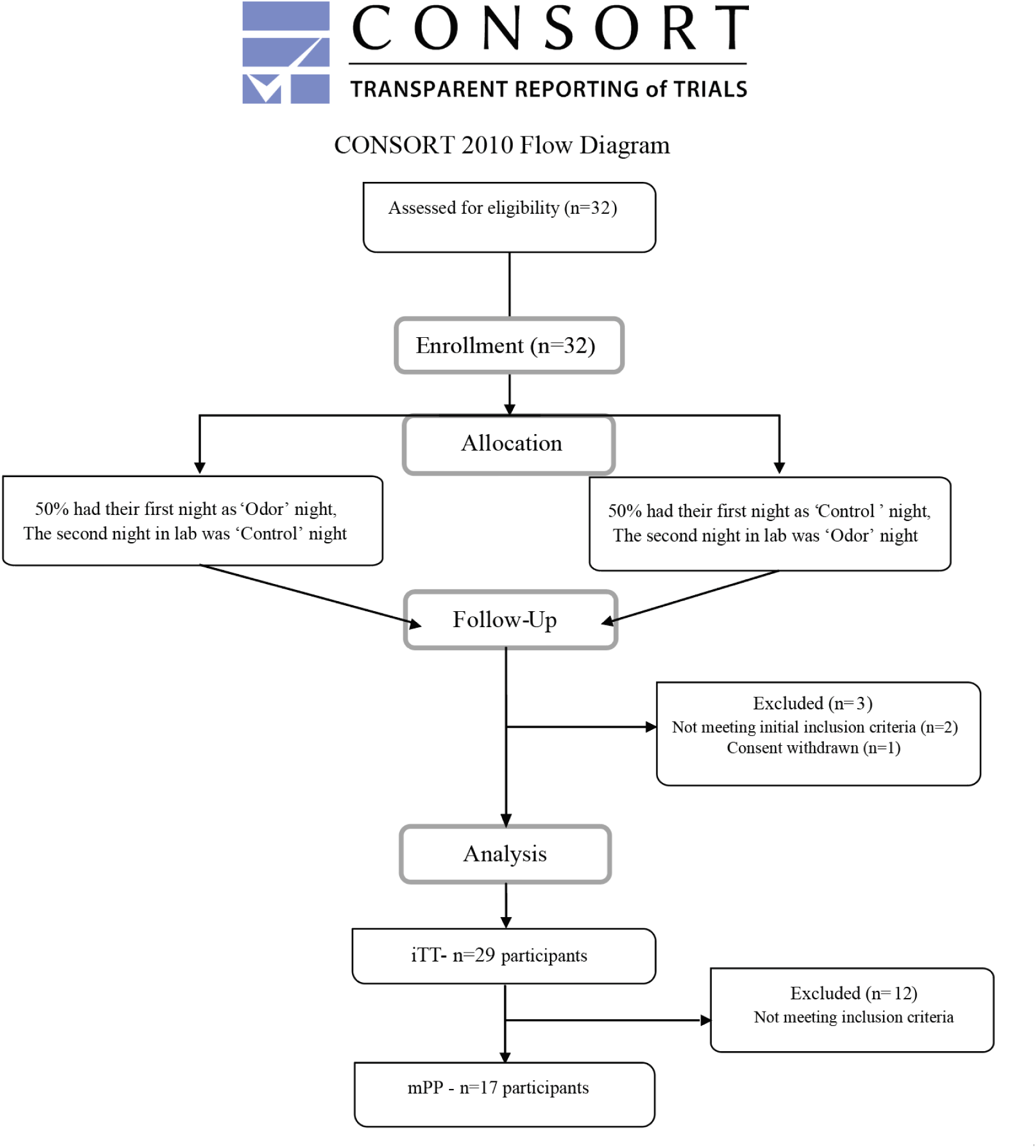
CONSORT diagram detailing the dynamics of patient enrollment and inclusion.

